# Generalisation of early learned tutor song preferences in female zebra finches (*Taeniopygia guttata*)

**DOI:** 10.1101/2022.05.05.490783

**Authors:** Jing Wei, Quanxiao Liu, Katharina Riebel

## Abstract

Song learning is a prime example for a culturally transmitted mating signal. Local or individual song variants are socially learned early in life and adults sing and prefer these songs. An unresolved issue in this context is the question of how learned preferences for specific variants generalise to songs sufficiently similar to the original model. Here we asked whether female zebra finches would generalise early learned song preferences along a similarity gradient based on syllables sharing between test and tutor songs. For each female, this gradient consisted of their tutor’s (father’s) song (F), two variants of unfamiliar songs edited to share 2/3 (F_2/3_) and 1/3 (F_1/3_) of syllables with father’s song and an unfamiliar song (UF). Females’ preferences were measured in a 4-way operant choice arena where the birds could perch on different operant perches to trigger playbacks of the four different songs. Number and duration of perch visits were positively associated with the number of syllables that the assigned stimuli shared with fathers’ songs. These results suggest that female zebra finches generalise early learned song preferences to songs sharing syllables (and/or voice characteristics) with songs learned early in life.

## 1. Introduction

Generalisation is the process whereby a learned response to a specific stimulus will also occur in response to a sufficiently similar novel stimulus and the intensity of response will be influenced by the degree of similarity between the novel and the initial stimulus (Purtle, 1973; Cheng, 2002; Ghirlanda and Enquist, 2003). Generalisation processes were initially studied mostly with discriminatory learning tasks in laboratory settings but investigations have increasingly shifted towards including species-specific biologically relevant learning contexts (Ghirlanda and Enquist, 2003). Examples for this approach are investigations into generalisation in filial imprinting for colour and shape (domestic chicken, *Gallus g. domesticus;* Bolhuis and Horn, 1992), mating colour preferences (guppies, *Poecilia reticulata;* Godin et al., 2005), locations of landmarks during foraging (honeybees, *Apis mellifera;* Cheng, 1999) and sexual imprinting on beak colour (zebra finches, *Taeniopygia guttata;* ten Cate et al., 2006). Given the fast-increasing documentations of learned mating preferences across vertebrate and invertebrate taxa (ten Cate and Vos, 1999; Verzijden et al., 2012; Montero et al., 2013; Dion et al., 2019), the question arises how and to which extent generalisation processes need to be considered in preference learning and other forms of experience-based mating preferences and choices (Ryan et al., 2003; Godin et al., 2005).

Song preferences are a primary driver of mate choice in many songbirds, emphasising the importance of understanding song preference learning (Riebel, 2003; White, 2004). While there is now sufficient experimental evidence to show the importance of early experiences on adult preferences (Riebel, 2003), little is known on how these preferences are generalised. More specifically, it is unclear whether learned song preferences of females only entail the specific songs heard early in life or whether and to which extent specific song memories generalise to similar songs. Pioneering studies on song preference learning suggest at least some generalisations from learned preferences for the father’s song and other structurally similar songs (Miller, 1979) and after cross-fostering for unknown variants of the respective subspecies song (Clayton, 1990) or, in the case of brood parasites, a new host species song (Payne et al., 2000). These studies are suggestive of generalisation, but they did not quantify the similarity of the test stimuli with the tutor song, raising the question as to whether song preference will indeed decrease or increase along a gradient of similarity. Creating a suitable gradient is challenging, because songs vary along several dimensions (frequency and amplitude modulations, and their timing) but individual singers can have repertoires of many variants of the species-specific song. In other species, however, such as the zebra finch, that sing only a single song type could be a suitable model to investigate this question. Moreover, as zebra finches are already an established model for song and song preference learning, the timing of the sensitive phase for song preference learning is well known (Riebel, 2003) and suitable song preference testing paradigms are available (Holveck and Riebel, 2007; Riebel, 2009). An additional advantage is that only males produce the elaborate courtship song in this species (although both sexes are very vocal throughout the day and seasons, Zann, 1996), meaning that song preference learning in females can be studied without the added confounds arising from perceptual biases arising from song production learning (Riebel et al., 2002; Gobes et al., 2009).

First evidence for early learned song preferences in female zebra finches stems from a series of phonotaxis experiments by Miller (1979). Females that could approach one of two loudspeakers preferred their father’s song over unfamiliar song. However, preferences for females’ fathers’ songs were weaker when they were tested against a structurally more similar song. As all subjects were raised by their biological fathers, within family sharing of perceptual biases could explain aligning song and song preferences, but subsequent studies using cross-fostering or tape tutoring to demonstrate preference learning were beyond doubt (reviewed in Riebel, 2003, 2009). Several of the earlier song preference studies reported observations suggestive of generalisation: preferences for learned tutor songs were weaker when paired with structurally more similar songs in phonotaxis tests (Miller, 1979; Clayton, 1988). However, male and female zebra finches that were tested with songs of unfamiliar brothers (from an earlier clutch of their parents), did not show a preference for the song of their unfamiliar brother over another unfamiliar song although, just like the test subjects, they had learned their song from the same father (Riebel and Smallegange, 2003). Perhaps effects were too weak to be picked up in this relatively small-scale study. However, song element overlap between similar songs was at least 75% in Clayton (1988) while the unfamiliar brothers in the study of Riebel and Smallegange (2003) shared only on average 53% ± 10% of the father’s song elements (unfamiliar songs showed 19% ± 11% elements random sharing with fathers’ songs). In a large-scale long-term breeding study, Wang et al. (2022) found that female zebra finches’ mating preferences in mixed aviary settings were best explained along the ‘micro-dialects’ of the separate rearing lines which females had experienced early in life. The combined results so far suggest that number of similar/learned elements/syllables in similar songs may affect the preference for early learned songs and that investigating the generalisation of learned song preference more systematically should in the first instance test a gradient of shared song elements/syllables.

In the present study, instead of relying on spectrograms based on human observer’s judgement of similarity of existing songs, we decided to manipulate the number of shared syllables as an approximation of a similarity gradient to increase stimulus control while keeping the acoustic properties and song structure of typical zebra finch songs. The song similarity gradient was defined by the proportion of syllables (1, 2/3, 1/3, 0) shared with the tutor and was created by exchanging syllables of the tutor song with syllables from an unfamiliar song. Stimulus sets consisting of these four variants of each female’s tutor’s song were then presented simultaneously in a circular 4-way operant setup to test whether female zebra finches would generalise their early learned song preferences for the tutor song to songs sharing syllables with the tutor song. We predicted that if females generalise their early learned preferences to songs sharing syllables with learned songs, their preferences should be the stronger the more syllables sharing there is between test and fathers’ songs.

## 2. Materials and methods

### 2.1 Subjects and housing

Subjects were the female offspring (N = 23) from 12 different breeding pairs from the zebra finch colony at Leiden University. All subjects had been raised by their biological parents in standard breeding cages (length × width × height: 80 × 40 × 40 cm) until day 65, which coincides with the end of the sensitive phase for song learning. In the breeding room, birds could not see adjacent neighbours but could see neighbours at a distance of ca. 3 m in cages placed along the wall on the opposite side of the room. In this setting, birds in our colony normally learn the song from the adult male they were housed with during the sensitive phase (here the father) in the same cage. From 65 dph (days post hatching) onwards, subjects were group housed in single-sex aviaries (group size: 20–25 birds/aviary, 175 × 80 × 200 cm) in a stock room with a total of 8 aviaries where zebra finch songs and calls were audible from within birds’ own and neighbouring aviaries. Birds were housed in these group aviaries until participating in preference testing at age 272–284 dph. Throughout, birds had *ad libitum* access to water, cuttlefish bone and commercial tropical seed mixture (Deli Nature 56-Foreign finches super, Schoten, Belgium) enriched with minerals and vitamins (GistoCal, Raalte, the Netherlands). In addition, germinated seeds, grated apple and/or carrot were provided twice a week and egg food was provided once a week. The housing rooms were kept at 20–22°C and 40–60% humidity and illuminated with artificial lights (Philips Master TL5 HO 49W/830) from 0700–2030 hours (13.5 light : 10.5 h dark) with a 15 min twilight phase with the light fading in and out at the beginning and the end of each day.

### 2.2 Recording of fathers’ songs and stimulus preparation

Prior to the experiments, the undirected songs of all fathers (N = 12) had been recorded. A recording session started by moving a single male into a wire mesh cage (100 × 60 × 30 cm) positioned at 75 cm height on a shelf within a sound attenuated room (height: 250 cm, width × length irregular quadrilateral: 106 × 158 × 304 × 387 cm). Birds were moved in in the afternoon to acclimate and then to stay overnight to catch the peak of singing activity the next morning. A microphone (Sennheiser MKH50-P48) mounted on a tripod (at a height of 1.5 m) was located 0.5 meter in front of the cage and connected to the hard disk of a computer (DELL OPTIPLEX 3020, soundcard: TASCAM UH-7000) running an automatic recording software (Ishmael, v. 1.0.2.0, settings: sampling frequency of 44.1 kHz, accuracy of 16 bits, file format: .wav). Zebra finch song is delivered in bouts of several songs. Each song starts with 2–10 repetitions of introductory notes followed by 1–8 repetitions of a male’s individual motif (the individual unique sequence of elements and syllables of each male; see Sossinka and Böhner, 1980; Zann 1996).

From all recorded songs of a given male, one song exemplar was randomly selected (using Random.org, for an example see appendix Fig. A.1). The selected songs had an average of 3.8 ± 1.4 motifs (from N = 12 fathers). Songs were high pass filtered at 425 Hz (PRAAT v. 6.0.28) and normalized (maximum amplitude of −1.0 dB using Audacity v. 2.3.1, function: normalized). From each of these songs, the predominant motif (defined as the motif with the most common syllable sequence) was selected by inspecting all motifs of the song using the combined spectrogram and waveform display in PRAAT (Spectrogram settings: 0–12 kHz, window length: 0.005 s, window shape: Gaussian, fft-size: 1000, 250 steps, dynamic range: 70 dB). Introductory notes were excluded unless they occurred not only before the first but also in the subsequent motifs within the same song.

To prepare the test stimuli, each father’s predominant motif was used (N = 12). For all these motifs, the duration, structure and position of all syllables were compared using the combined spectrogram and waveform display in PRAAT. The mean duration of fathers’ motifs was 809 ± 171 ms (range: 520– 1080) with on average of 6.6 ± 1.4 syllables (range: 4–9). These motifs were paired into 6 stimulus sets of two motifs with comparable duration and syllable number (mean difference between pairs: 59 ± 40 ms, 0–2 syllables; for all details see appendix Table A.1). In the experimental design, each stimulus set was then assigned to test the daughters of the two males that had contributed the two songs in a given stimulus set. In this way, each male’s song motif was the familiar song motif (F) for his daughter(s) as well as the unfamiliar song motif (UF) for the daughters of the other paired male and vice versa (see an example in Fig. 1). For each father’s motif, two additional variants labelled F_2/3_ and F_1/3_ were edited by replacing 1/3 and 2/3 syllables of one motif with syllables from the paired motif in the set (see Fig. 1).

**Fig. 1.**
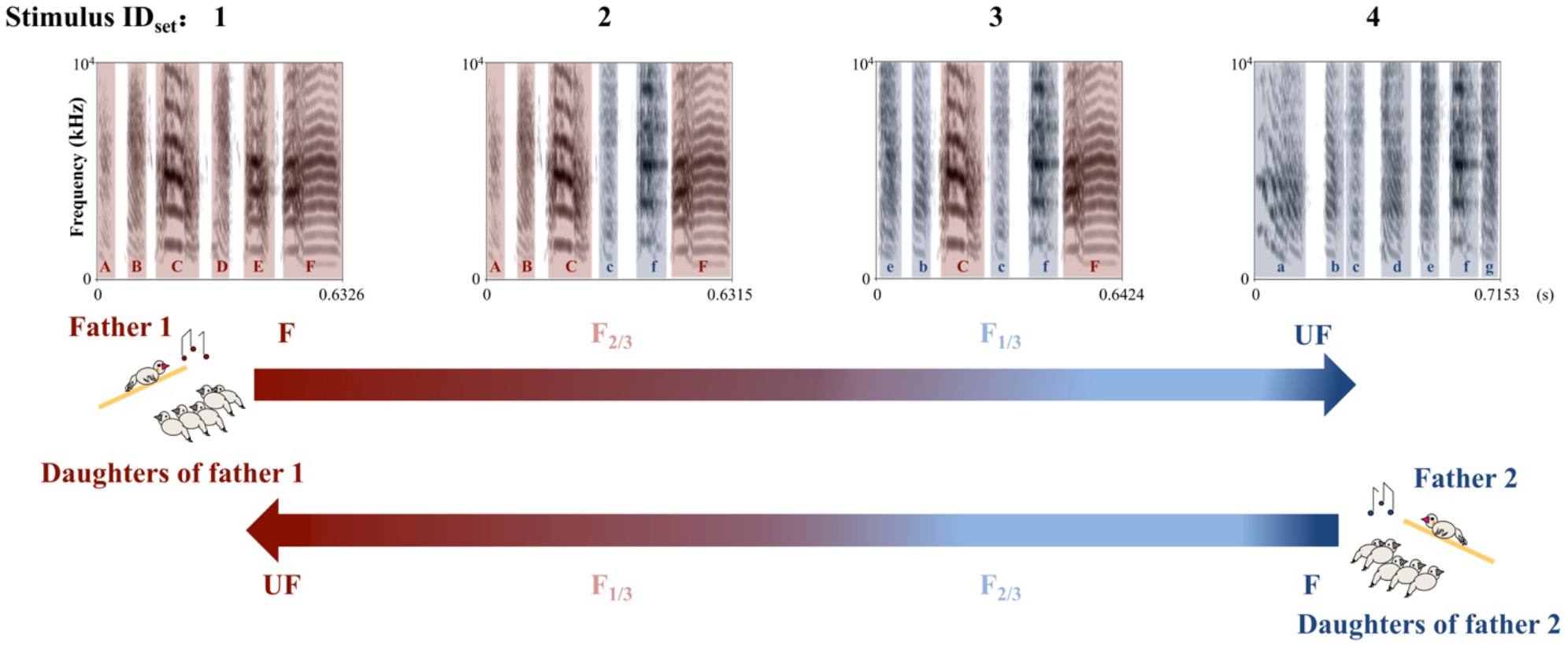
Example of a stimulus set consisting of four song stimuli (ID_set_: 1, 2, 3, 4). Song 1 and 4: song motifs of fathers 1 and 2 were used as familiar song for their daughters and as unfamiliar song for the daughters of the other male; Song 2 and 3 (F_2/3_ and F_1/3_): unfamiliar song motif with 2/3 (F_2/3_) and 1/3 (F_1/3_) syllables of the father’s song motif, and 1/3 (F_2/3_) and 2/3 (F_1/3_) syllables of the unfamiliar song motif. The colour indicates from which male the syllables originated and the colour gradient in the arrows below the spectrograms illustrate the gradient of shared syllables from F (dark red, syllables A-F) to UF (dark blue, syllables a-g). NB: only one motif per stimulus category is shown, but the stimuli used in the test consist of a song with several motif repetitions (for examples of complete stimulus set see appendix Fig. A.1).

For pairing of motifs and subsequent syllable exchange, we used the combined spectrogram and waveform display and the Annotate-TextGrid function of the PRAAT software (Spectrogram settings: 0–12 kHz, windows length: 0.005 s, dynamic range: 70 dB) to measure and label all syllables within each set of the paired motifs. All syllables (excluding the subsequent silent pause) were then parsed into separate sound files.

Syllables within motifs paired in the design were ranked by duration, structure and position (for a detailed description of the editing procedure, see supplementary materials). Syllables matched for similarity (2/3 or 1/3 in each song rounding up to the nearest integer) were exchanged between the songs of the two fathers to yield either the F_2/3_ or F_1/3_ stimulus by copying and pasting the syllables one by one. The newly edited motifs did not differ on average more than 16 ± 4 ms (2.1% ± 1.5%) from the original familiar (F) motif (range: 0–58 ms (0–10.4%)). The father’s motif and unfamiliar motif were control edited and underwent the same dissembling steps but were then reassembled in the original order (for an example of a stimulus set, see appendix Fig. A.1). All edited motifs were digitally multiplied with the original number of motif repetitions in the original songs.

### 2.3 Apparatus

Most song preference tests offer two choices only (Riebel et al., 2002; Holveck and Riebel, 2010) but an earlier operant study in our lab had already successfully tested female zebra finches’ song preferences offering four song choices via four different operant buttons placed in the back panel of an experimental cage (Ritschard et al., 2010). In this setup the four pecking keys were aligned in one row and although females used all four pecking keys, they preferred the two buttons at the outer two perches over the two buttons near the central two perches. For the present study, we therefore used a different setup: a modification of an 8-way mate choice arena (henceforth referred to as the ‘carrousel’, see ten Cate et al., 2006; Holveck et al., 2011) which because of its circular shape allowed us to present four song stimuli equidistant and equally reachable from the centre of the arena to prevent edge effects (see Fig. 2).

**Fig. 2.**
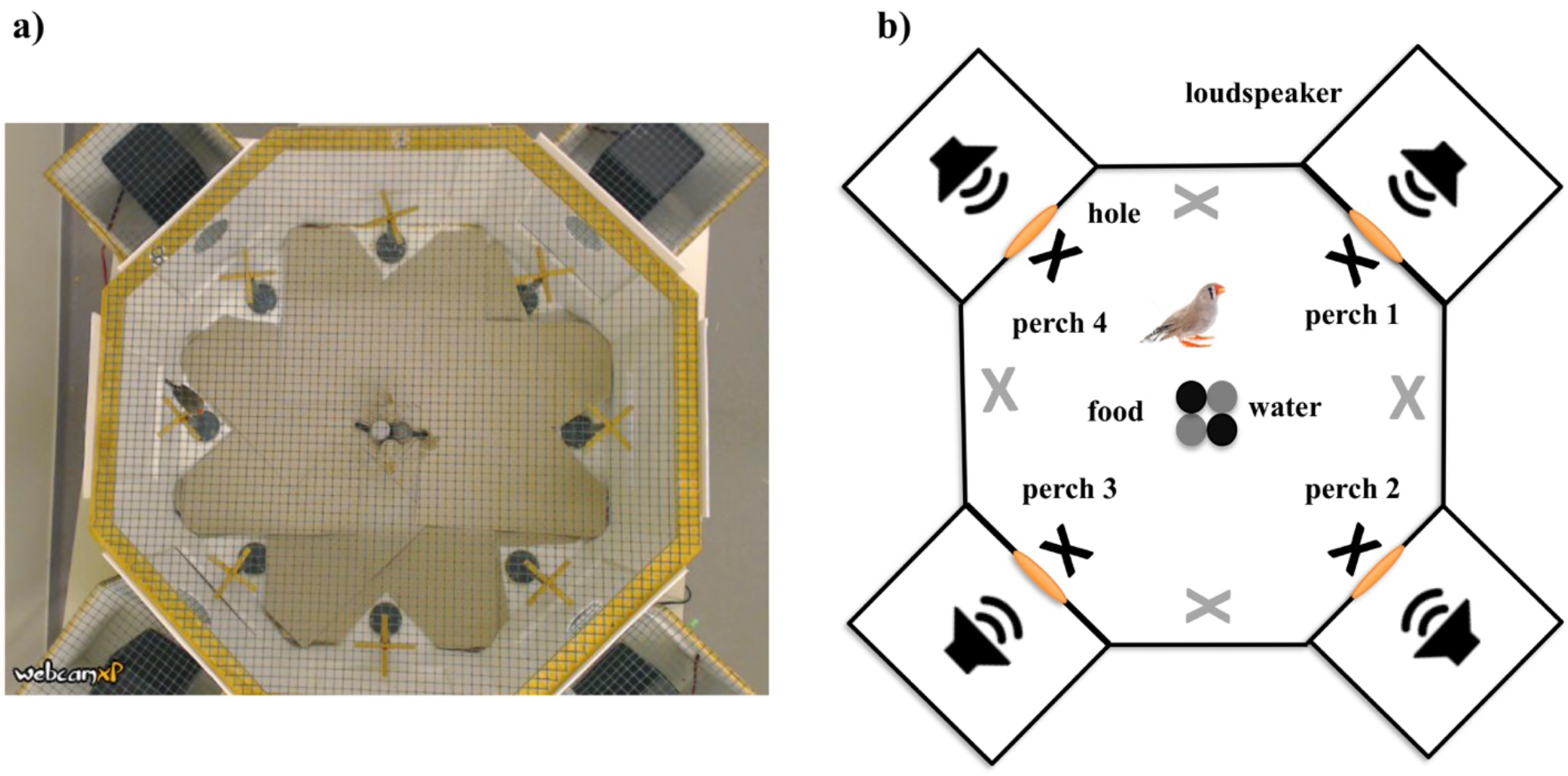
The testing apparatus: a) Photograph directly above the apparatus taken in the test room. b) Top view of the apparatus. The round openings on the white boards in front of the loudspeakers are indicated in orange colour. The black crosses indicate the operant perches (1–4) and the grey crosses the control perches. Landing on an operant perch triggered a single playback of the stimulus song assigned to it (F, F_2/3_, F_1/3_, UF). Song stimuli associated with the four operant perches were rotated one position clockwise every two hours. Two food and water dispensers were placed in the middle of the carrousel (marked with black and grey solid circles).

The carrousel had a top covering of wire mesh (height: 35 cm, diameter: 70 cm) and 8 arms with small wire mesh openings to 8 small removable cages (length × width × height: 26 × 26 × 35 cm). For the experiment described here, all arm were closed off with white plastic board dividers, but every other arm contained a small loudspeaker (CB4500, Blaupunkt, Hildesheim, Germany) that could broadcast via a round opening (diameter = 3 cm) in the central part of the plastic dividers (see Fig. 2). Birds could trigger song playbacks by landing on the small cross shaped perches in front of the loudspeaker openings. This triggered a microswitch connected to a computer placed outside the experimental room that controlled the playbacks and registered the moment the bird landed and departed (Dell OptiPlex 3010 with RADEON HD soundcard and custom written software in Visual basic 6, by P.C. Snelderwaard). Hopping on an operant perch always immediately triggered a single song playback no matter how long a subject stayed on the perch but if a subject left the perch before the end of the song playback to fly to another operant perch, the playback stopped immediately and the new playback associated with the new perch was triggered. Playback levels on each perch had been set to maximum level of 68 dBA (re: 20 μPa) prior to the test (VOLTCRAFT SL-100 sound meter, settings: sound level high, A-weighting).

The experimental setup was placed on a table (height: 75 cm) in a sound attenuated chamber (height: 250 cm, width × length irregular quadrilateral: 335 × 280 × 290 × 300 cm). To reduce acclimation time (Waas et al., 2005; Adrian et al., 2022), an additional loudspeaker (JBL Clip2, Vietnam) was placed on the floor directly under the carrousel broadcasting a continuous recording of one of our bird rooms (45 dBA sound peak pressure level measured at the height of the perches inside the carrousel). A camera (Logitech HD 1080p, Lausanne, Switzerland) was mounted 1.5 meter above the carrousel and connected to the computer (software: Webcam 7 pro, v. 1.5.3.0) controlling the setup and allowing to monitor females’ behaviour during trials from outside the experimental room.

### 2.4 Preference test

To start a trial, the playback of colony sound via the loudspeaker under the carrousel, the operant setup and the camera above the apparatus were switched on before moving the focal subject into the setup between 0900–0930. Each of the four song stimuli was assigned to one of the perches (starting with assigning F, F_2/3_, F_1/3_ and UF to perches 1, 2, 3 and 4; for perch location see Fig. 2b). Every two hours (called one ‘block’), the stimuli assigned to particular perches were rotated one position clockwise until each stimulus had been presented at each position once. The whole test with four blocks lasted 8 hours (ending between 1700–1730). While all birds had the same start figuration, the different run-in times (i.e. time to start showing the operant behaivour) differed among females. This means that females encountered different perch*song stimuli combinations when they first started to discover the perches. All females were tested again 2–3 months (79 ± 11 days) after the first test with the same stimuli to test whether their song preferences were repeatable (as observed for the operant setup involving key pecks (Riebel, 2000; Holveck and Riebel, 2010)) as this setup previously had only been used for association tests with live stimulus birds (ten Cate et al., 2006; Holveck et al., 2011) but not as an operant setup.

### 2.5 Ethical note

We adhered to the ASAB/ABS Guidelines for the Use of Animals in Research (ASAB, 2021) and the European and Dutch legislation on animal experimentation. Birds were housed socially apart from the 8 hours preference test. Behavioural observation in an open choice arena is not considered a procedure in the Experiments on Animals Act (Wod, 2014) which is the applicable legislation in the Netherlands in accordance with the European guidelines (EU directive no. 2010/63/EU) regarding the protection of animals used for scientific purposes. At all times, all birds were housed and cared for in accordance with these regulations and internal guidelines concerning care of the animals by licensed and skilled personnel and all procedures were reviewed and monitored by the official Animal Welfare Body responsible for monitoring and implanting legal requirements. Throughout testing, food and water were available *ad libitum* and after testing, females were returned to their home aviaries.

### 2.6 Data Analysis

The registration software logged a time stamped location ID for each landing and departure from an operant perch (original file format: text, transferred and saved into excel spreadsheets, v. 16.33). From these data, the number of perch visits and duration of each perch visit (per location and per stimulus) were extracted. For song preference analyses, only females that had visited all song stimuli at least once were included (as preference strength across stimuli can only be assessed meaningfully after a female had encountered all choices). Additionally, after visiting all stimuli at least once, females also had to remain active and show at least 40 perch visits (see appendix Table A.2).

Data were analysed using the statistical software R (functions lmer and glmer in package lme4 in R 4.1.2). To test whether females had repeatable preferences, we used a generalized liner mixed-effects model with perch visits per stimulus category (per block) in the 2^nd^ test round as the response variable and perch visits in the 1^st^ test round as a covariate (Model 1, see Table 1). Stimulus category and block were fixed factors (ordered), female’s ID and stimulus set were included as two random effects (intercepts).

**Table 1.** Repeatability of females’ preferences in the first and second preference test. The table shows the results of a generalized linear mixed-effects model analysis of perch visits per stimulus category in each block during the two preference tests two months apart for the subsample of females that met the test criteria in both tests.

| Model | Chisq | Df | p value |
| --- | --- | --- | --- |
| Model 1: Perch visits in the 2<sup>nd</sup> test |  |  |  |
| Stimulus category | 44.13 | 3 | < 0.001 |
| Block | 53.75 | 3 | < 0.001 |
| Perch visits in the 1<sup>st</sup> test (covariate) | 636.34 | 1 | < 0.001 |

*GLMM with a Poisson distribution. Female's ID and stimulus set were included as two random intercepts (N = 10 females).*
*Marginal and conditional R<sup>2</sup> for this model are 0.30 and 0.94.*

The four song stimuli were modelled as stimulus categories F, F_2/3_, F_1/3_, and UF, but for some analyses the individual songs within a stimulus set were also modelled as factor stimulus ID_set_ with the four levels 1, 2, 3 or 4 (see example in Fig. 1 and further explanation in appendix Table A.3). This was because using stimulus category did not allow to uniquely identify a specific song within a set because each tutor song was the familiar song for some females but the unfamiliar song for other females (see Fig. 1). To test whether some stimulus songs were accidentally more attractive than others (Tchernichovski et al., 2021), we first checked whether stimulus ID_set_ within a stimulus set influenced females’ preference (by modelling the number of perch visits as the response variable, stimulus ID_set_ as a fixed factor, block as an ordered factor, and female’s ID and stimulus set as two random effects (Model A1, see appendix Table A.4). We then ran this model again but now instead of stimulus ID_set_, with stimulus category as response variable (ordered: F, F_2/3_, F_1/3_, UF; Model 2, see appendix Table A.4). We then compared these two models (Model A1 with Model 2) with ANOVA (see appendix Table A.4) and kept the model with the lower AIC. For this final model, we computed estimated marginal means (R package ‘emmeans’, ‘emmeans’ function) to see how levels within the fixed factors affected the response variable (see appendix Table A.5). As the data registration in the carrousel also allowed for measuring the visit durations for each song stimulus, we used the same procedure to test how stimulus ID_set_ and stimulus category affected visit duration but replaced perch visits with duration per visit (Log-transformed) as the new response variable in two linear mixed-effects models (Models A2 & 3, see appendix Tables A.4 & A.5).

## 3. Results

Of the 23 females that participated in the first test, 6 females did not visit all song stimuli during the 8-h test and one female showed fewer than 40 perch visits (for individual results, see appendix Table A.2), These 7 females were excluded from further analysis. The remaining 16 (70%) females had visited all song stimuli, and had been very active (with 661 ± 26 perch visits, range: 143–1926) within the 8-h period; in comparison the excluded females visited the operant perches only 14 ± 4 times (range: 0–78) during this period. Females which reached the criteria had visited all song stimuli at least once after 138 ± 96 min (range: 31–300) from the start of testing. In the second test two months later, activity was 442 ± 21 perch visits (range: 80–2046) and only 11 of 23 females (i.e. 48%) reached the criteria (of those, 10 had met the criteria in the first test round, see appendix Table A.2). Comparing the preference of the 10 females that had been included in both tests showed their preferences for the four song stimuli were repeatable (Model 1, see Table 1), i.e. their perch visits to the four different stimulus categories in the first test round predicted visiting behaviour in the second test round (see appendix Fig. A.2). Given that preferences were repeatable, but that the number of responsive females in the second test round was so much lower than in the first test round, all following analyses were conducted on the data of the 16 females from the first test round only (as there were sufficient females per stimulus set to compare daughters from both males that provided songs for the F/UF song sets).

Looking at females’ behaviour in the first test round in more detail revealed that females differed in how fast they discovered all perches (see appendix Fig. A.3), but by the end of the second block, 75% (12/16) females had visited all four operant perches and had heard all four song stimuli. The majority of females (14/16) started to hop on the perches during the first block (0–2 h) and the other 2 females (F712 and F758) visited the first perch during the second block (2–4 h). The cumulative individual learning curves (see appendix Fig. A.3) showed a slow increase in the operant behaviour changing into a steep increase, which is typical of learning curves (Pearce, 2008). Notably, when looking at the cumulative plots (see Fig. 3) showing the number of females that had visited a particular stimulus category per unit time, from the four stimulus categories separately, the father’s song seemed to have been discovered faster. However, as there had been no indication to females which perch coded for which song, this likely means that once females had (accidentally) discovered the familiar father’s song they were slower to explore the remaining perches.

**Fig. 3.**
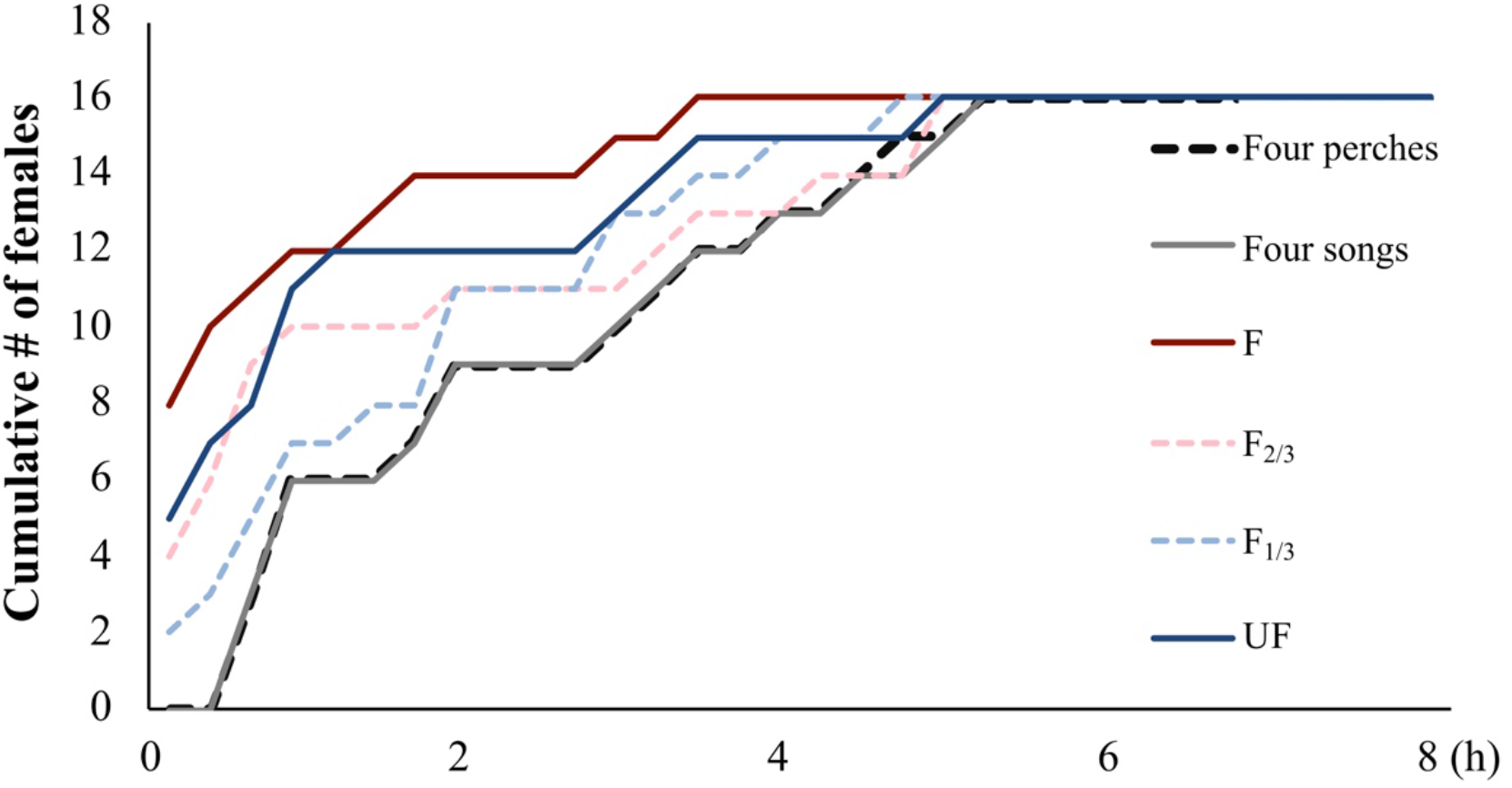
Cumulative frequencies of the number of females that had discovered a particular stimulus category, all perches or all four songs. In each curve counts are added cumulatively over time: a count is added each time a female had visited a specific perch, stimulus category or all perches or all songs for the first time. F: father’s song; UF: unfamiliar song; F_2/3_ and F_1/3_: two edited versions of the unfamiliar song with 2/3 and 1/3 of the syllables originating from the father’s song.

For the 16 females that had met the criteria in the first test, we investigated their preferences for songs along the similarity gradient in two steps. We first visually inspected the frequencies with which females chose the four song categories and then inspected the average visit duration of each female for each stimulus category. Fig. 4a illustrates the predictions arising from generalisation which should lead to a linear decrease of the number of perch visits along the similarity gradient; the expected pattern if there were learned preferences for the fathers’ songs but without generalisation (Fig. 4b) or if there was an absence of preference which should lead to random (roughly equally high) numbers of visits to all perches (Fig. 4c). The observed number of perch visits and the average duration per visit per stimulus category for the tested females (after birds visited all song stimuli once) were shown in Fig. 4d and 4e. Note that across the six tested stimulus sets, the highest number of visits were in the majority of cases observed for the fathers’ songs (the red symbols for song 1 or 4 in each set, see Fig. 4d & e). Statistical analyses (see below) of the data showed that visits both for the father’s song and for the remaining song categories were not random.

**Fig. 4.**
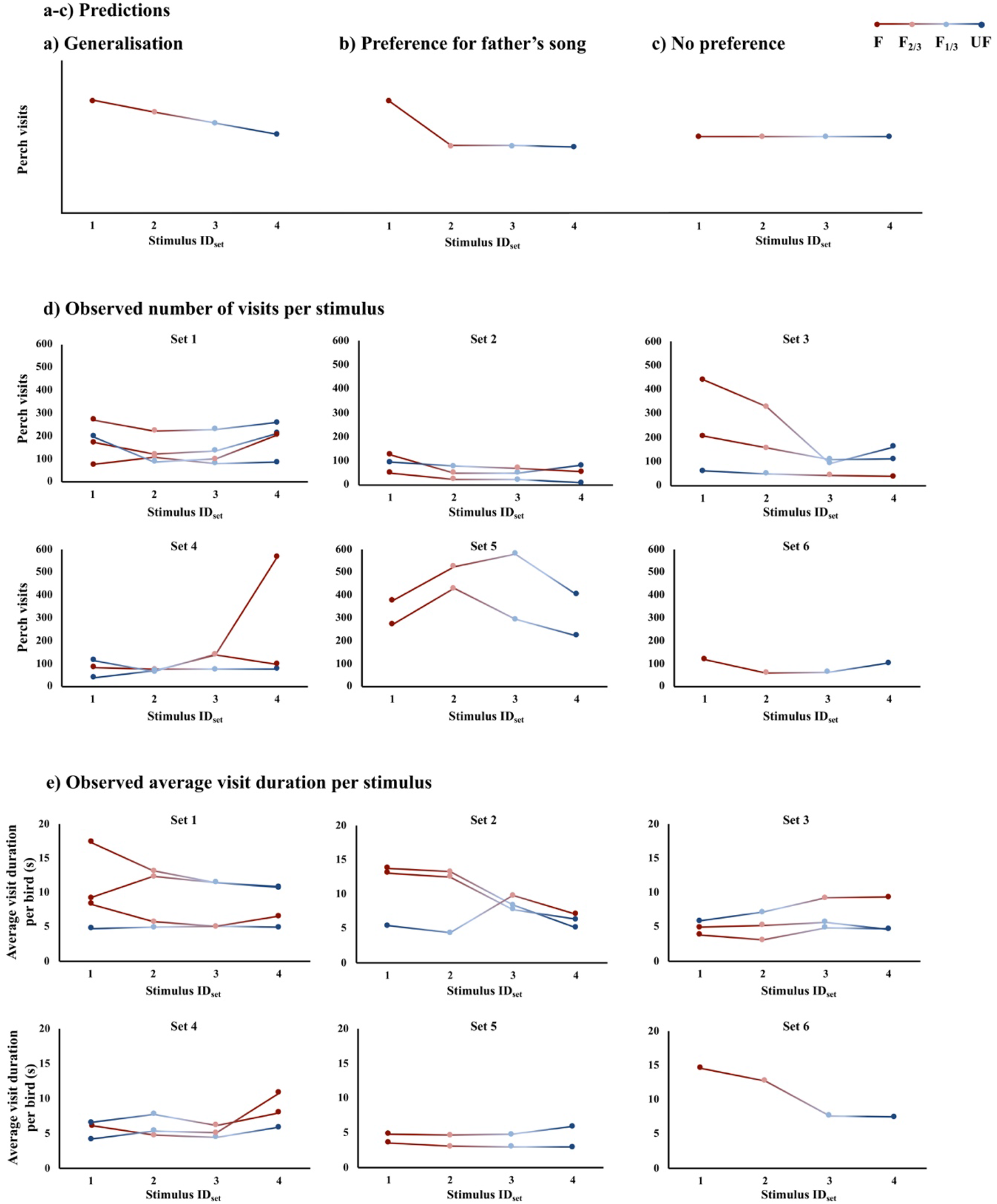
Expected and observed perch visits and visit duration for the different song categories along the similarity gradient. a) Expected preference with generalisation, b) preference for father’s song but without generalisation (ID_set_: 1, 2, 3, 4) and c) no preference. d) Observed number of perch visits and e) average visit duration per bird for each stimulus category (ID_set_: 1, 2, 3, 4) in each stimulus set (1–6) of 16 females after they had visited all song stimuli once. Each dot in d) refers to the total number of perch visits on that stimulus category and in e) presents the average visit duration of each bird for that stimulus category. The colours of the symbols and lines code the four stimulus categories (F: dark red, F_2/3_: pink, F_1/3_: light blue, UF: dark blue).

Across all birds, the number of birds’ perch visits and visit duration on each stimulus category were better explained by stimulus categories (see Fig. 5 and Table 2) than the factor stimulus ID_set_ (ΔAIC = 255.3 and 97.1, see appendix Table A.4). Perch visits were found to be affected by stimulus category (see Fig. 5a, Model 2 in Table 2 and appendix Table A.5): the fathers’ (F) songs were visited more often than the other three categories. Perch visits to F_2/3_ songs were higher than to both F_1/3_ and UF songs, and perch visits to F_1/3_ songs were comparable to UF songs. The visit duration was also found to be affected by stimulus category (see Fig. 5b, Model 3 in Table 2 and appendix Table A.5): F songs elicited longer duration per visit than all other categories. Visit duration to F_2/3_ songs was comparable to F_1/3_ songs but was longer than UF songs. F_1/3_ songs were comparable to UF songs. In general, the higher the percentage of familiar syllables, the more often birds visited and the longer the birds stayed on the perch for those songs. There was also a time effect independent of stimulus category that the longer the test progressed, the more frequently females visit the perches and the shorter they stayed on each perch (see Models 2 & 3 in appendix Table A.5).

**Fig. 5.**
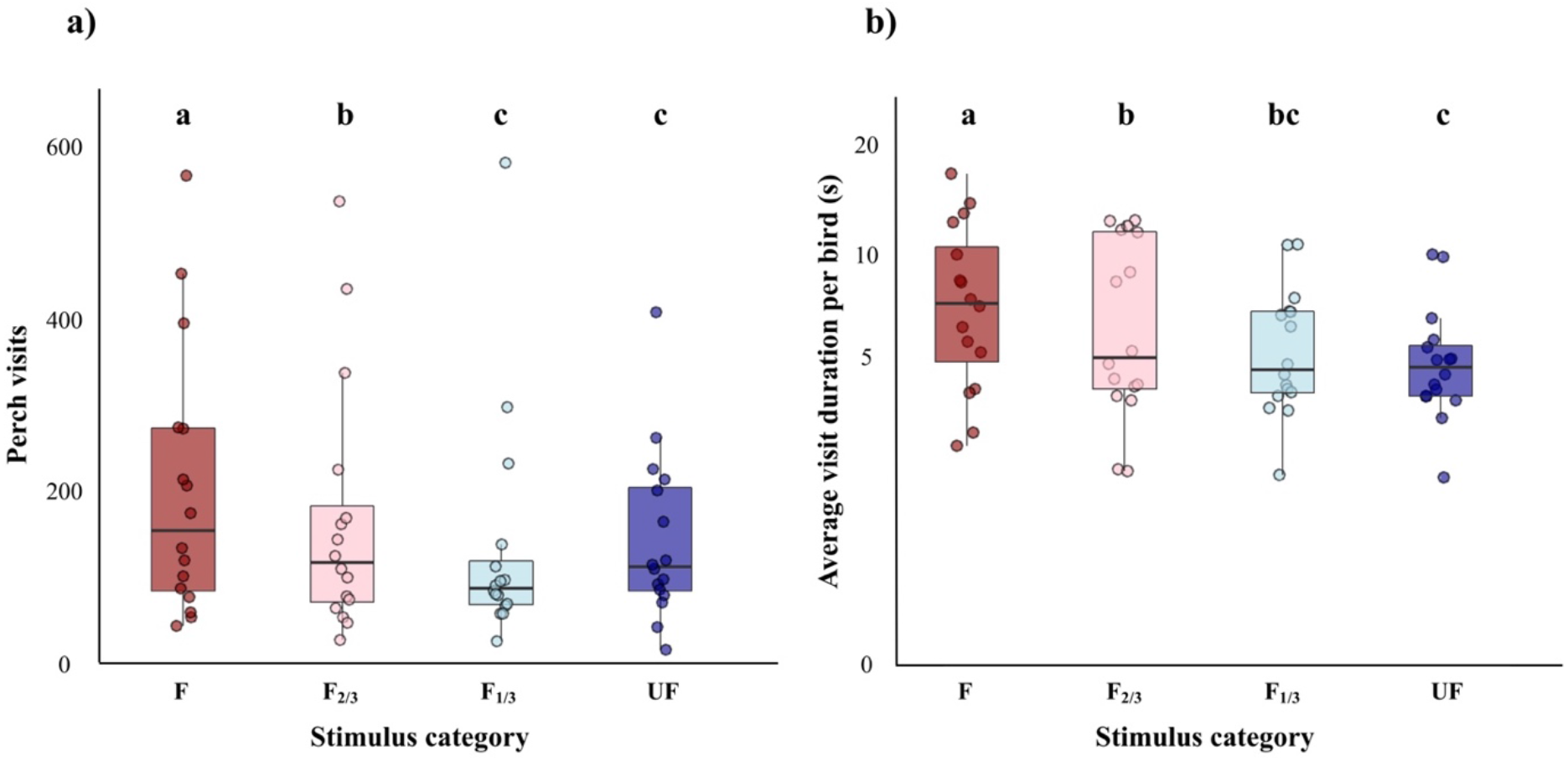
a) Perch visits per stimulus category. Box plots show the median, first and third quartiles. Whiskers show the 1.5 times interquartile range. The dots show the absolute number of visits per bird per song stimulus category. b) Each female’s average visit duration per stimulus category. Different letters in a) and b) show significant differences between groups (p < 0.05 emmeans contrast, for details see Models 2 & 3 in appendix Table A.5). F = father’s song, UF = unfamiliar song, and F_2/3_ and F_1/3_: songs blending 2/3 and 1/3 of the father’s song syllables with syllables from the unfamiliar song.

**Table 2.** Effects of stimulus category and test block on female zebra finches’ preferences (generalized linear mixed-effects and linear mixed-effects models).

| Model | Chisq | Df | *p* value |
| --- | --- | --- | --- |
| Model 2<sup>a</sup>: Perch visits |  |  |  |
| Stimulus category | 265.21 | 3 | < 0.001 |
| Block | 854.39 | 3 | < 0.001 |
| Model 3<sup>b</sup>: Visit duration |  |  |  |
| Stimulus category | 104.02 | 3 | < 0.001 |
| Block | 419.58 | 3 | < 0.001 |

<sup>a</sup> Generalized: Poisson distribution; <sup>b</sup> Normal distribution. Female's ID and stimulus set were included as random intercepts. Marginal and conditional *R*<sup>2</sup> are 0.30 and 0.96 for Model 2, and 0.05 and 0.27 for Model 3. The estimates for block as covariate in Models 2 and 3 are 0.26 and -0.19 respectively (see appendix Table A.5).

## 4. Discussion

Male and female songbirds learn about family and population specific songs early in life from conspecific tutors and these early song templates shape adult song and song preferences. In this study, we investigated whether female zebra finches generalised learned song preferences for their tutor’s song along a gradient of songs edited to share decreasing number of syllables with the originally learned song. Overall, female zebra finches preferred their tutor’s (here their father’s) song which is according to expectation arising from earlier studies in this species (reviewed in Riebel, 2003, 2009). Across the test song gradient, the number of syllables shared with the father’s song affected females’ preferences: the more familiar tutor song syllables there were in the songs, the more and the longer birds visited the perches triggering playbacks of these songs. This behaviour is in line with predictions arising from generalisation of early learned song preferences for songs sharing syllable and/or voice characteristics with the tutor song.

Notably, females’ early learned preferences for their father’s song were repeatable when tested again two to three months later, just as previously reported for preferences for early learned songs in operant key pecking and live male preference tests (Riebel, 2000; Holveck and Riebel, 2010; Holveck et al., 2011). This suggests that using operant perches and four song stimuli simultaneously is a suitable method to test a stimulus gradient. A song preference test involving a similarity gradient has not previously been conducted in songbirds (but see for a test of generalisation with peak shift for a visually imprinted beak colour preference in ten Cate et al. (2006)). Females’ preference strength increased with the number of syllables originating from the preferred (father’s) song. This finding allows reinterpretation of previous results that were suggestive of generalisation of learned song preferences in zebra finches (Miller, 1979; Clayton, 1988) but which had not tested multiple options along a similarity gradient. These studies observed that (learned) preferences for fathers’ or tutors’ songs were weaker if tested against structurally more similar than more dissimilar songs (females in Miller, 1979; young males and females in Clayton, 1988) but no such effect was observed when males or females were tested with songs of their biological, but unfamiliar brothers (Riebel and Smallegange, 2003). Albeit that the songs of these birds shared significantly more elements with the tutor (father) than unfamiliar control songs (unfamiliar brothers’ songs: 53% ± 10%; unfamiliar songs: 19% ± 11%), they were not preferred. This is surprising given that other studies in the zebra finch reported preferences for (foster) brothers’ songs and peer group songs (Holveck and Riebel, 2010; Honarmand et al., 2015) or microdialects (Wang et al, 2022). However, sample sizes in Riebel and Smallegange (2003) were small overall (18 birds from only four families) so that the different learning outcomes in tutees from different tutors (Tchernichovski et al., 2021) could have had an unduly influence. Even small differences in performance among singers with songs learned from the same tutor can influence song (Holveck and Riebel, 2010) and live male preferences (Holveck et al., 2011), the overall picture arising from all these studies is thus that early learning does result in preferences for specific songs but also songs resembling these songs (Riebel 2003, 2009; Wang et al., 2022).

On the subspecies level, generalisation of learned song preferences has already been shown to be of consequence for pair formation: cross-fostering between the two subspecies of zebra finches (*Taeniopygia g. guttata* and *Taeniopygia g. castanotis*) made females prefer structurally different song of their rearing, rather than parental subspecies (Clayton, 1990). On a within-population level, after just a few generations of separate breeding of several lines in the laboratory, females’ mating preferences in mixed aviary settings were best explained along the ‘micro-dialects’ of the rearing lines that females had experienced early in life (Wang et al., 2022). In an experimental approach, Le Maguer et al. (2021) raised females with different artificial dialects. When tested in operant song preference tests, these females preferred (unfamiliar) songs from their rearing dialect (Le Maguer et al., 2021). These studies demonstrate that early experience can change preferences and mating decisions and thus quickly affect realised pairings (see also how song preference learning might lead to new host species in brood parasites, Payne et al., 2000). An important point to consider is that all the above studies used natural songs from different birds to test preferences whereas in the current experiments we blended syllables of the songs of two singers. As zebra finches can identify individuals by voice characteristics (Elie and Theunissen, 2018; Geberzahn et al., 2021), and songs in our study shared familiar (or preferred) syllables directly taken from the tutor song (i.e. the gradient was both about shared structural features and shared voice characteristics), it is feasible that either or both structural features of song units of individual voice characteristics were recognisable for zebra finches and guided their preferences. However, overall, the experimental and observational evidence of our current study and the different studies discussed above are supporting the notion that zebra finches show some generalisations of their early learned songs, for which the current study provides experimentally support by using for the first time a gradient of song similarity in an experimental situation. Note that our paired song motifs were biased towards a greater similarity than two randomly selected songs of the same population would be. However, the results from different cross-fostering studies show that young females generalise from their foster subspecies (Clayton, 1990) or their foster micro dialects (Wang et al., 2022) or artificial dialects (Le Maguer et al., 2021).

Albeit that generalisation of learned song preferences has not been studied systematically to date, the observed generalisation ties in with results in other songbirds. While we tested within-population variation, a number of studies have shown generalisation processes to be at play on the subspecies and between species level. Cross-fostered female zebra finches showed preferences for the specific songs of males as the same subspecies as their foster fathers (Clayton, 1990). Female indigobirds (*Vidua chalybeate*) imprinted on a novel foster species and as adults showed preferences for the conspecific males that mimicked the songs of the new host species (Payne et al., 2000). Female song sparrows (*Melospiza melodia*) showed stronger preferences for playback of strangers’ songs that were more similar to their mate’s song (O’Loghlen and Beecher, 1999). Female swamp sparrows (*Melospiza georgiana georgiana*) tutored with particular song from their local population as adults showed preferences for unfamiliar local song over unfamiliar foreign song suggesting the possibility of the generalisation of features in learned songs and the application of these features when discriminating among novel songs in adulthood (Anderson et al., 2014). The combined evidence suggests that generalisation of learned preference influences the process of mate choice based on song variation on the within-population level and that these processes need to be further investigated in the future.

The question of how learned mating preferences are generalised is of relevance not only for songbirds, but a wide variety of animal taxa, and insights from discirmination learning tasks in the laboratory are informative for understanding the potential consequences of such processes. In many species, mating preferences are acquired during a learning process which is referred to sexual imprinting where juvenile experiences with social models – often the caregiver(s) will shape adult mating preferences (reviewed in ten Cate and Bateson, 1988). A behavioural response bias called ‘peak shift’ can arise from discrimination learning, meaning that animals have a stronger response to novel stimuli away from a positively rewarded stimulus in a direction opposite from a negatively or neutrally rewarded stimulus, and vice versa (Weary et al., 1993; ten Cate and Rowe, 2007). Learned mating preference in the visual domain have already demonstrated peak shift in songbirds. Male zebra finches raised with mothers with artificial orange or red beaks, when presented with a gradient of beak colours ranging from more extreme on the paternal to more extreme on the maternal side as adults, chose a colour hue shifted away from the imprinted hue, providing evidence for generalisation and peak shift (ten Cate et al., 2006). Learned acoustic and visual mating preferences have also been found to affect mate choice in other taxa and modalities (Bolhuis and van Kampen, 1992; Dukas, 2004, 2005; Dion et al., 2019 for reviews). For example, female guppies (*Poecilia reticulata*) can learn mating preference for particular males from other females and then generalise these learned mating preference to other males with a similar colour phenotype (Godin et al., 2005). In the vocal learning domain, peak shift has been demonstrated in zebra finches in the responses to novel songs with higher or lower numbers of odd elements (i.e. along the intensity gradient) than the training stimuli (Verzijden et al., 2007). Geberzahn and Derégnaucourt (2020) found that both male and female zebra finches could correctly classify novel stimuli from trained stimuli which were learned from the same tutor in the generalisation phase, suggesting that they are able to discriminate the individual unique song type and they were also shown to pay more attention to syllable structure than syllable order. The increasing documentation of generalisation in the context of learned mating preferences therefore calls for more research into its functional significance.

In conclusion, the findings reported here provide experimental evidence that female zebra finches generalise their early learned preferences for a specific song to novel songs sharing more or fewer syllables with the specific song. These results can provide a point of departure for additional, more systematic tests of generalisation of learned preferences for acoustic mating signals and their consequences for mate recognition and selection in songbirds. On a methodological note, this study also suggests that operant song preference tests can be expanded from classic 2-way paradigms to include at least 4-way choices. The modification of the mate choice arena (‘carrousel’) into an operant setup proved suitable to test females’ preferences among four different song playback options simultaneously and might possibly use with even more options in the future, thus allowing to test larger, more complex and perhaps more realistic options sets (many individuals with similar songs) in the future. This could greatly aid to further investigate generalisation of early learned song preferences by offering more detailed gradients of shared syllables and to further test whether shared structural or shared voice characteristics are driving the generalisation of preference.

## Acknowledgement

We would like to thank Carel ten Cate for very helpful comments on this manuscript. We also thank Peter Snelderwaard for technical support and Michelle Geers for caring for the birds. This work was supported by Institute of Biology, Leiden University. Jing Wei and Quanxiao Liu were funded by the Chinese Scholarship Council.

## Appendix A. Supplementary data

### Editing rules for exchanging syllables from two paired song motifs

As zebra finches prefer long songs (Vyas et al., 2009), we always chose the two fathers’ song motifs that were the most similar in duration (0.809 ± 0.171 s) for the same stimulus set. The selection of syllables that were to be exchanged between two motifs in the same set occurred according to the following three rules: 1) syllables were ranked by duration - if there were several syllables of the same duration then 2) those closest in song structure and 3) in the most similar position were chosen. Editing occurred as follows: two paired motifs were opened in the PRAAT software using the combined spectrogram and waveform display and the Annotate-TextGrid function (Spectrogram settings: 0–12 kHz, windows length: 0.005 s, dynamic range: 70 dB) to measure and label all syllables. Syllables in each set of two paired motifs were then ranked by duration, structure and position. The two syllables in paired motifs that were of the most similar duration were first selected to be exchanged. We replaced the syllable in the familiar song motif with the syllable from the unfamiliar song motif by using the copying and pasting functions in PRAAT. If there were two or more syllables of similar duration that could be exchanged, we compared syllable structure and then exchanged the two most similar syllables. If there were several syllables of similar structure, the syllable which was closest in sequential position was chosen. This was repeated until 1/3 or 2/3 of syllables were exchanged thus resulting in the F_2/3_ or F_1/3_ stimuli.

**Fig. A.1.**
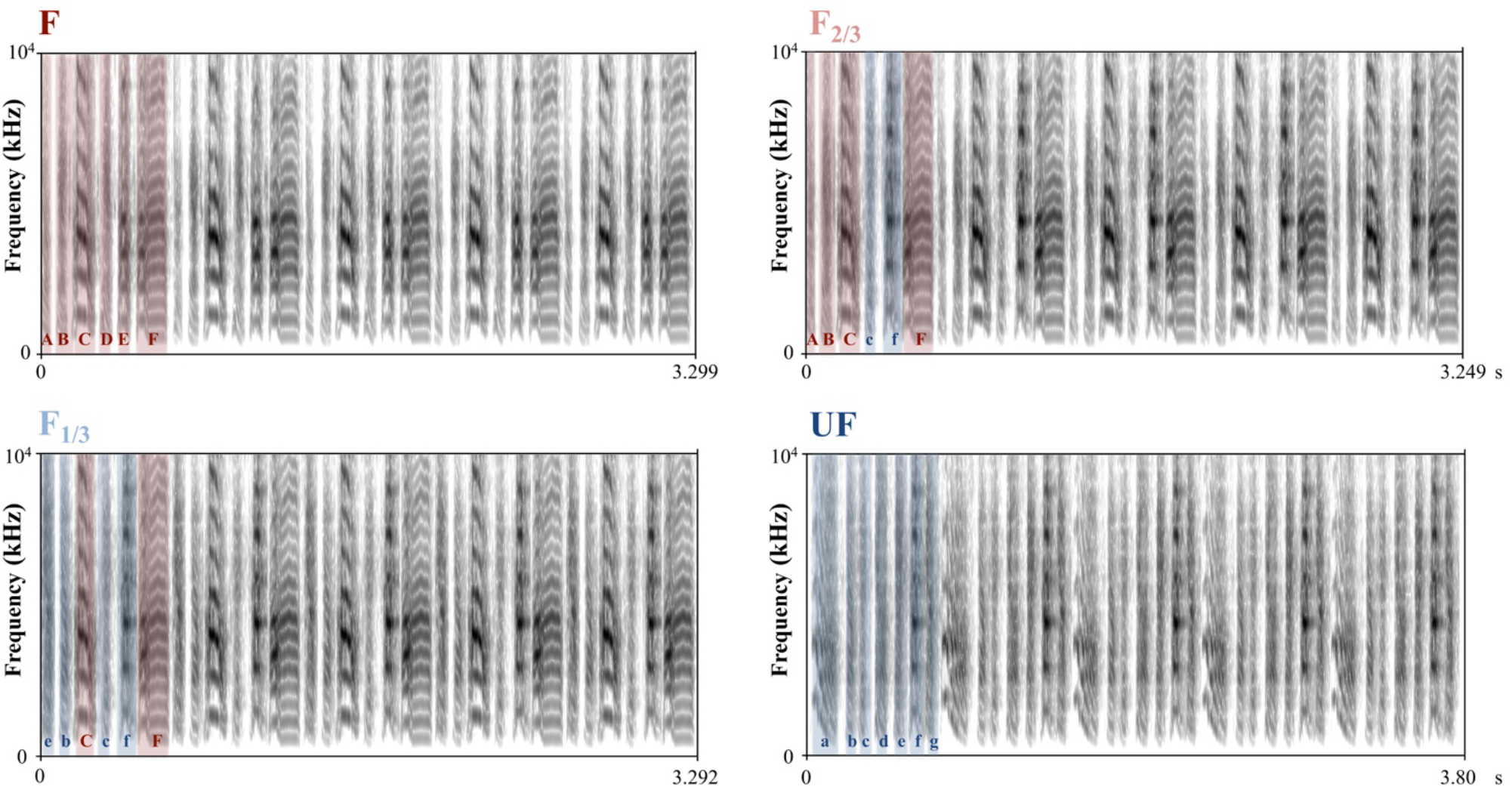
Example of a complete stimulus set consisting of four song stimuli (the complete songs with several motif repetitions correspond to the four single motifs in Fig. 1. F: father’s song; F_2/3_: father’s song with 2/3 of syllables remaining and 1/3 of syllables exchanged; F_1/3_: father’s song with 2/3 of syllables exchanged; UF: unfamiliar song. The colour highlighting of the syllables in the first motif indicate from which male the syllables originated.

**Table A.1.**
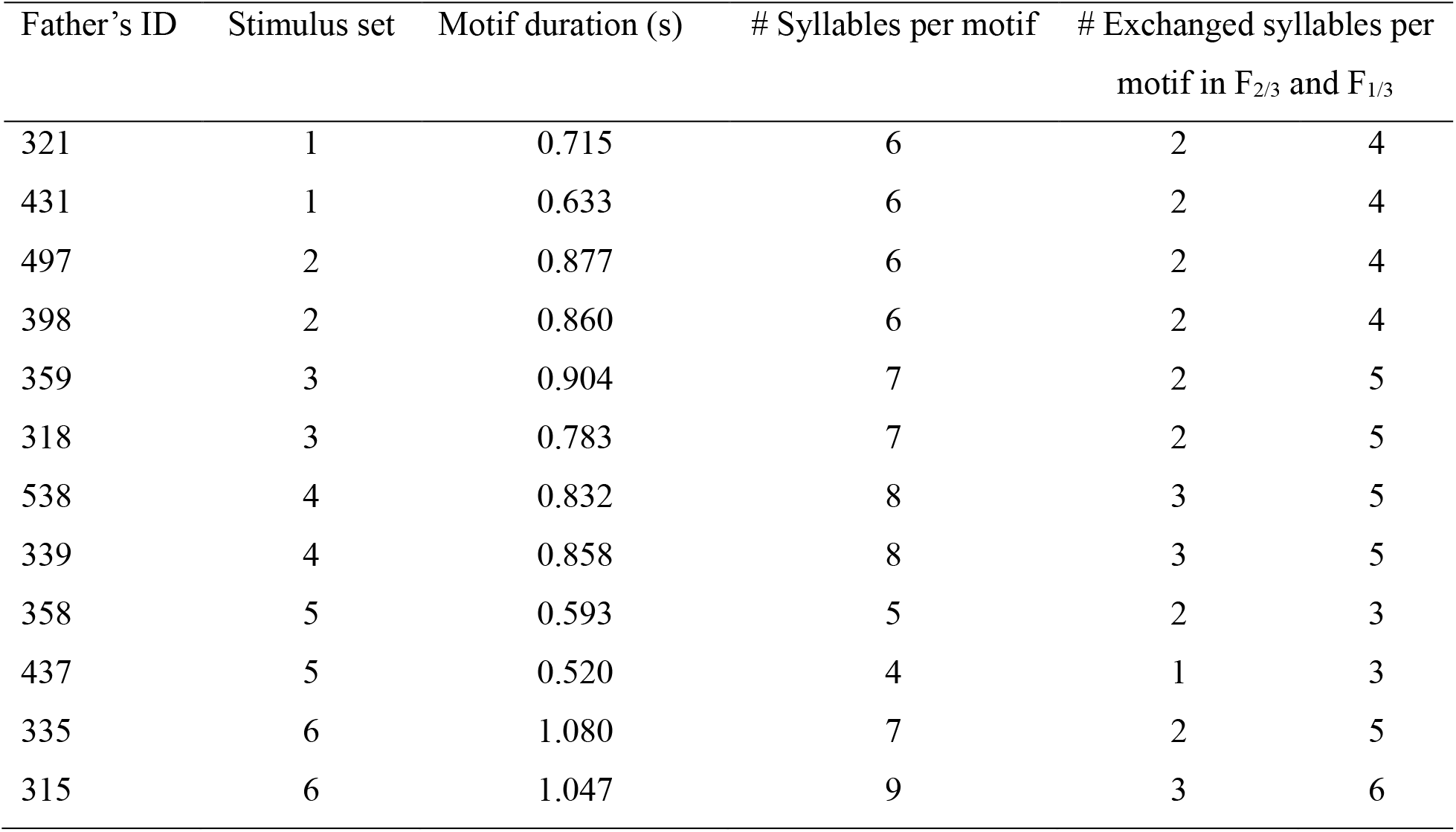
Father’s ID, stimulus set, motif duration, the number of syllables per motif and exchanged syllables per motif in F_2/3_ and F_1/3_.

**Table A.2.** Individual females’ perch visits to the different song stimuli (after having reached criterion of visiting all stimuli at least once) and at which block they had first visited all stimuli at least once.

| Female's ID | Stimulus set |  | Test round | Perch visits |  |  |  | Total visits | All stimuli visited > 1 at block# |
| --- | --- | --- | --- | --- | --- | --- | --- | --- | --- |
|  | F | UF |  | F | F<sub>2/3</sub> | F<sub>1/3</sub> | UF |  |  |
| 720 | 321 | 431 | 1 | 172 | 122 | 135 | 211 | 640 | 1 |
|  |  |  | 2 | 0 | 0 | 0 | 0 | 0 | - |
| 721 | 321 | 431 | 1 | 270 | 222 | 229 | 259 | 980 | 1 |
|  |  |  | 2 | 201 | 164 | 113 | 109 | 588 | 1 |
| 723 | 321 | 431 | 1 | 74 | 108 | 81 | 85 | 348 | 2 |
|  |  |  | 2 | 0 | 0 | 0 | 0 | 0 | - |
| 730 | 431 | 321 | 1 | 204 | 97 | 88 | 198 | 587 | 2 |
|  |  |  | 2 | 33 | 26 | 31 | 30 | 120 | 1 |
| 712 | 497 | 398 | 1 | 51 | 24 | 23 | 13 | 111 | 3 |
|  |  |  | 2 | 39 | 64 | 106 | 105 | 314 | 1 |
| 715 | 497 | 398 | 1 | 131 | 51 | 55 | 89 | 326 | 1 |
|  |  |  | 2 | 2 | 2 | 2 | 3 | 9 | 4 |
| 760 | 398 | 497 | 1 | 56 | 71 | 78 | 95 | 300 | 3 |
|  |  |  | 2 | 36 | 43 | 56 | 12 | 147 | 4 |
| 761 | 398 | 497 | 1 | 0 | 0 | 0 | 0 | 0 | - |
|  |  |  | 2 | 2 | 3 | 4 | 3 | 12 | 4 |
| 732 | 359 | 318 | 1 | 450 | 335 | 94 | 162 | 1041 | 1 |
|  |  |  | 2 | 4 | 7 | 0 | 0 | 11 | 4 |
| 733 | 359 | 318 | 1 | 211 | 159 | 110 | 113 | 593 | 1 |
|  |  |  | 2 | 79 | 37 | 9 | 22 | 147 | 3 |
| 754 | 318 | 359 | 1 | 41 | 44 | 55 | 68 | 208 | 2 |
|  |  |  | 2 | 49 | 36 | 12 | 17 | 114 | 2 |
| 753 | 318 | 359 | 1 | 0 | 0 | 0 | 0 | 0 | - |
|  |  |  | 2 | 20 | 15 | 8 | 30 | 74 | 2 |
| 743 | 538 | 339 | 1 | 84 | 75 | 76 | 76 | 311 | 2 |
|  |  |  | 2 | 24 | 48 | 43 | 35 | 150 | 2 |
| 742 | 538 | 339 | 1 | 7 | 22 | 1 | 3 | 33 | 3 |
|  |  |  | 2 | 19 | 11 | 5 | 2 | 37 | 2 |
| 756 | 339 | 538 | 1 | 564 | 141 | 66 | 117 | 888 | 1 |
|  |  |  | 2 | 202 | 89 | 62 | 148 | 501 | 1 |
| 758 | 339 | 538 | 1 | 98 | 166 | 93 | 39 | 396 | 3 |
|  |  |  | 2 | 0 | 0 | 0 | 0 | 0 | - |
| 745 | 358 | 437 | 1 | 271 | 432 | 295 | 223 | 1221 | 1 |
|  |  |  | 2 | 101 | 84 | 93 | 75 | 353 | 1 |
| 748 | 358 | 437 | 1 | 392 | 534 | 579 | 405 | 1910 | 1 |
|  |  |  | 2 | 531 | 619 | 479 | 409 | 2038 | 1 |
| 770 | 437 | 358 | 1 | 0 | 0 | 0 | 0 | 0 | - |
|  |  |  | 2 | 0 | 0 | 0 | 0 | 0 | - |
| 718 | 335 | 315 | 1 | 117 | 61 | 64 | 107 | 349 | 1 |
|  |  |  | 2 | 0 | 0 | 0 | 0 | 0 | - |
| 717 | 335 | 315 | 1 | 0 | 0 | 0 | 0 | 0 | - |
|  |  |  | 2 | 0 | 0 | 0 | 0 | 0 | - |
| 728 | 315 | 335 | 1 | 0 | 0 | 0 | 0 | 0 | - |
|  |  |  | 2 | 0 | 0 | 0 | 0 | 0 | - |
| 729 | 315 | 335 | 1 | 0 | 0 | 0 | 0 | 0 | - |
|  |  |  | 2 | 0 | 0 | 0 | 0 | 0 | - |

**Table A.3.**
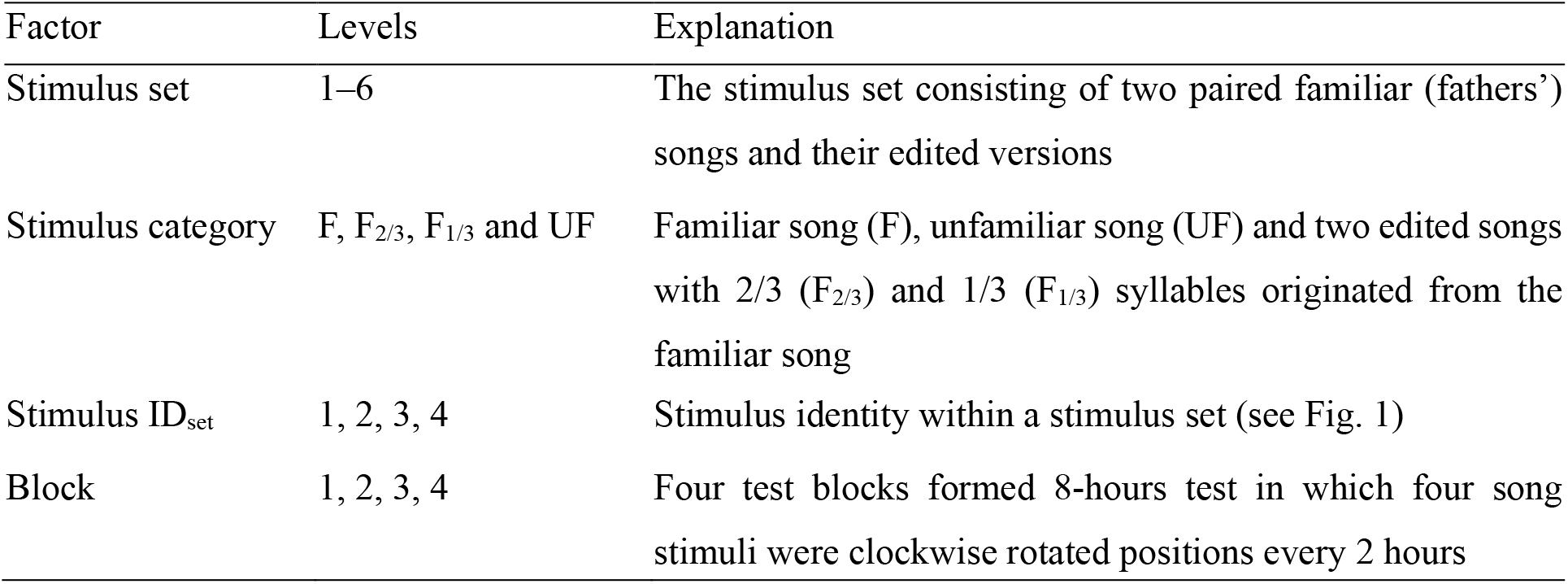
Fixed and random factors in linear mixed-effects and generalized linear mixed-effects models (Models 1–3, A1 & A2).

| Factor | Levels | Explanation |
| --- | --- | --- |
| Stimulus set | 1–6 | The stimulus set consisting of two paired familiar (fathers') songs and their edited versions |
| Stimulus category | F, F<sub>2/3</sub>, F<sub>1/3</sub> and UF | Familiar song (F), unfamiliar song (UF) and two edited songs with 2/3 (F<sub>2/3</sub>) and 1/3 (F<sub>1/3</sub>) syllables originated from the familiar song |
| Stimulus ID<sub>set</sub> | 1, 2, 3, 4 | Stimulus identity within a stimulus set (see Fig. 1) |
| Block | 1, 2, 3, 4 | Four test blocks formed 8-hours test in which four song stimuli were clockwise rotated positions every 2 hours |

**Fig. A.2.**
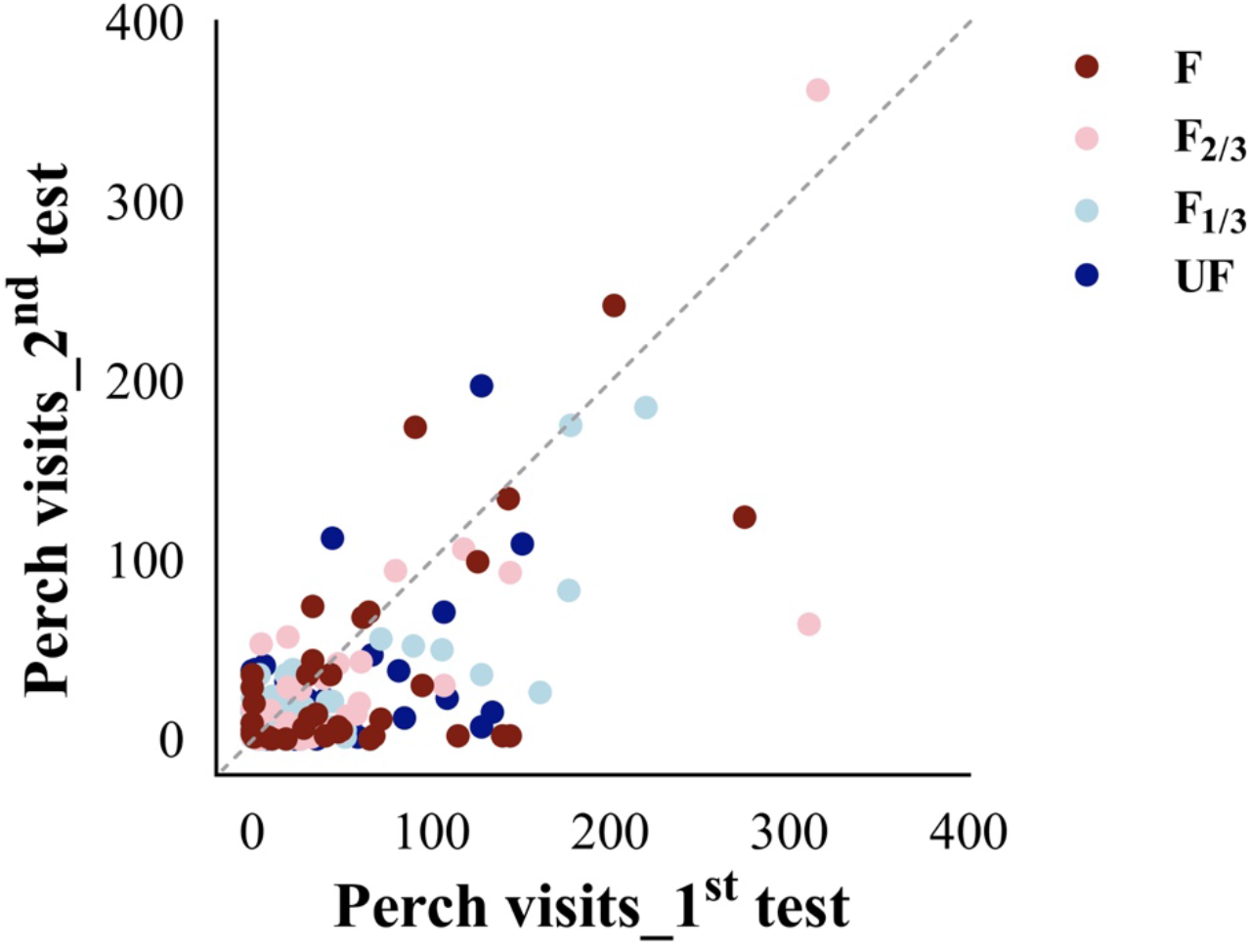
Perch visits to each song category in the 1^st^ and 2^nd^ test. Each point represents one bird’s perch visits to each stimulus category in each block. The four song stimulus categories (F, F_2/3_, F_1/3_, UF) are indicated by colours (dark red, pink, light blue, dark blue). The statistical analyses controlled for repeated measures within an individual (within and between tests), see Model 1 in Table 1.

**Fig. A.3.**
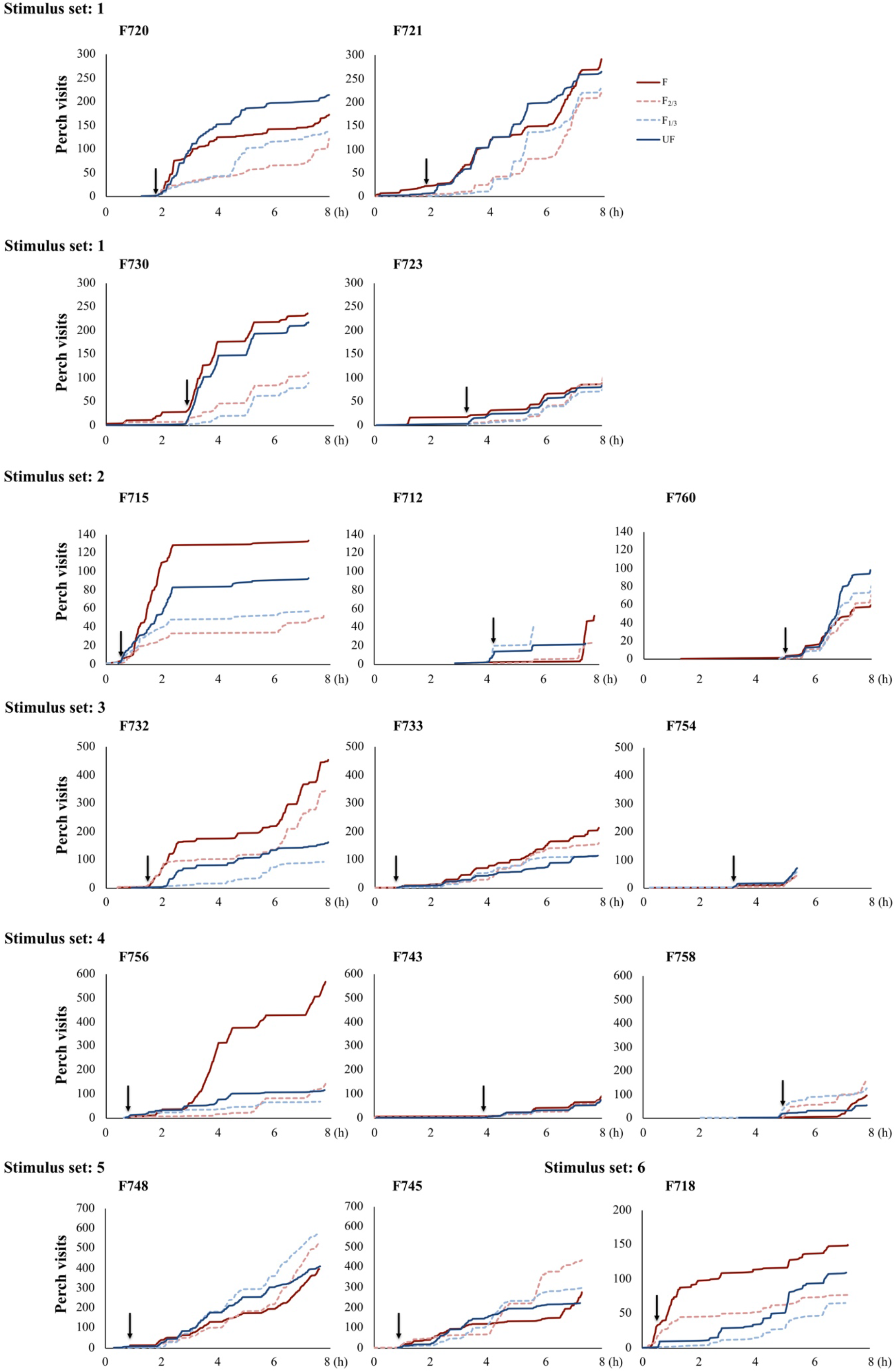
Cumulative plots showing individual females’ (N =16) perch visits to the four different stimulus categories. F: father’s song (dark red line); F_2/3_: father’s song with 2/3 of syllables remaining and 1/3 of syllables exchanged (pink dashed line); F_1/3_: father’s song with 2/3 of syllables exchanged (light blue dashed line); UF: unfamiliar song (dark blue line). Each plot shows all perch visits during an 8-h test. The black arrow for each female points the moment when a female had visited all song stimuli. Birds’ IDs and their stimulus sets’ numbers are shown above plots.

**Table A.4.**
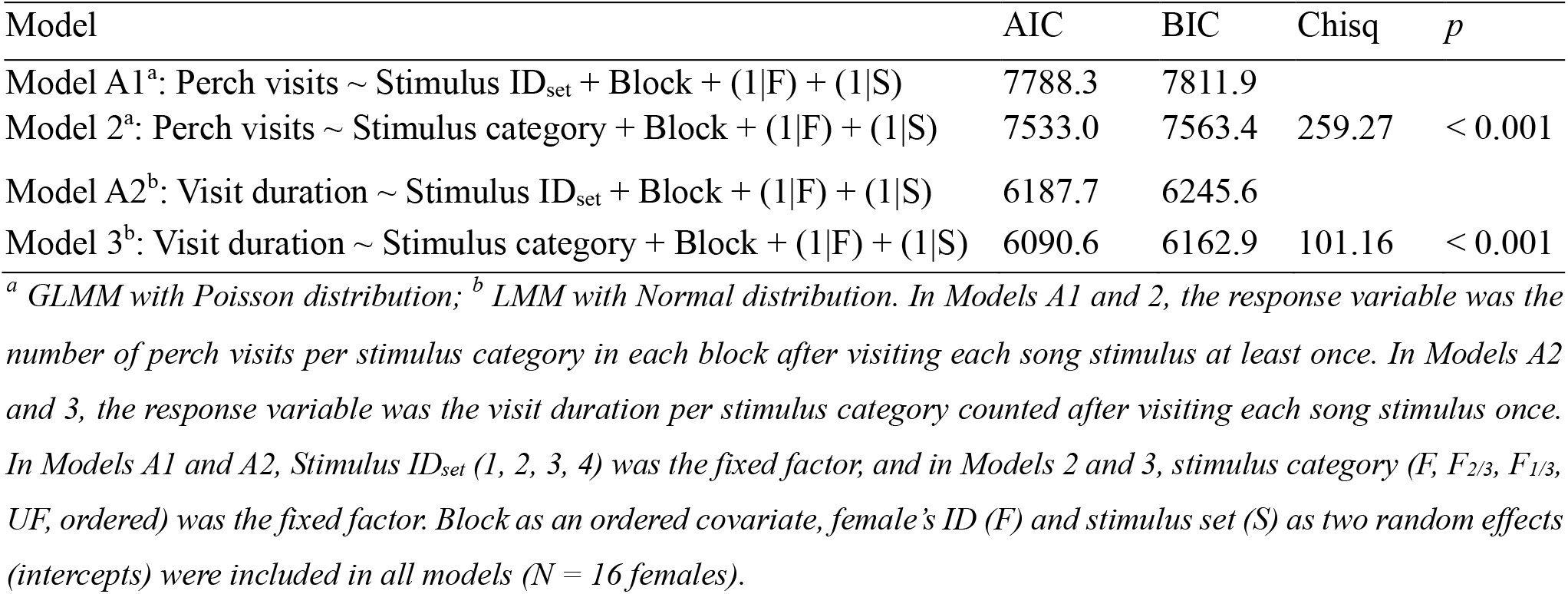
Results of model comparisons (Model A1 with Model 2, Model A2 with Model 3) in ANOVA tests.

**Table A.5.**
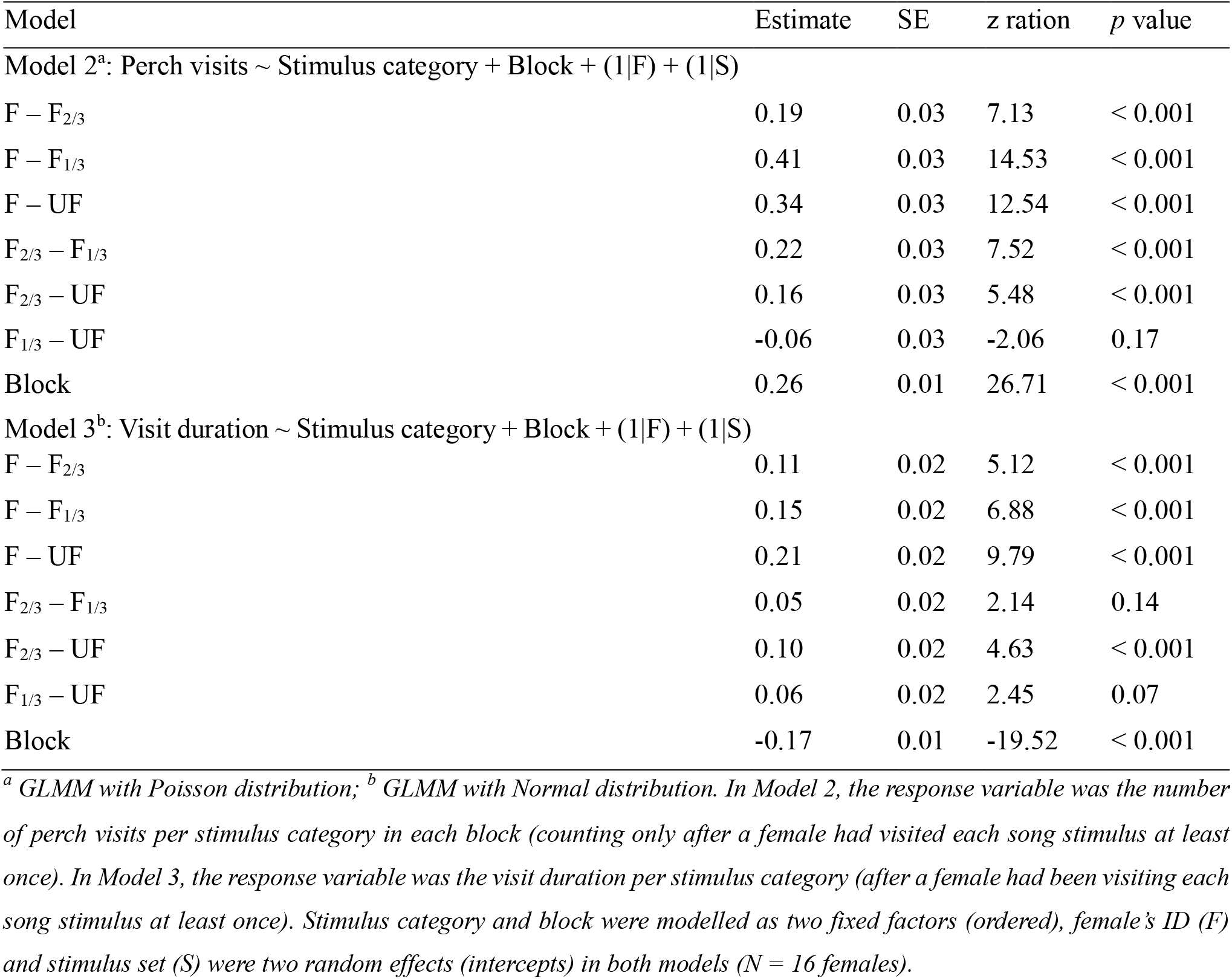
Effects of stimulus category and test block on female zebra finches’ preference (generalized linear mixed-effects and linear mixed-effects models, ordered factors: ‘stimulus category’, ‘block’). Contrasts (R package ‘emmeans’, ‘emmeans’ function) for each level within the fixed factors of Model 2 and Model 3 (see Table 2).

